# Neural Oscillations Coordinate Continuous Error Correction During Force Control

**DOI:** 10.1101/2025.08.07.669031

**Authors:** N. Menghi, E. Balestrieri, D. Grignolio, G. Coricelli, C. Hickey

## Abstract

Effective motor control depends on the brain’s ability to monitor performance and make continuous corrections. While many studies focus on discrete errors, everyday actions often require ongoing feedback-based adjustments. Here, we used an isometric force control task with EEG to investigate the neural dynamics supporting real-time error correction. Participants maintained a constant grip force with or without continuous visual feedback. With feedback, behavior showed ∼6 Hz rhythmic fluctuations, consistent with active correction. These fluctuations were mirrored in EEG activity across theta, beta, and alpha bands—oscillations linked to performance monitoring, updating, and attentional control. Without feedback, performance decayed linearly, and the corresponding neural signatures were reduced. These findings suggest that continuous sensory feedback engages a dynamic feedback loop involving distinct neural processes that support adaptive behavior. Our results highlight the importance of oscillatory activity in tracking and correcting moment-to-moment fluctuations in force, offering insight into the neural basis of feedback-loop force control.

## 1 Introduction

Motor control and error correction are fundamental processes that enable humans to produce precise and adaptive actions. For example, when we are cooking scrambled eggs, we must constantly adjust the temperature to avoid overcooking or undercooking. If the pan gets too hot, the eggs start to stick to the pan, prompting us to lower the temperature. If it cools down too much, cooking slows and so we raise the temperature again. This ongoing regulation depends on visual, tactile, olfactory, and auditory feedback to stay within the narrow temperature range that ensures the desired consistency. Hence, sensory feedback provides continuous updates about our actions and performance accuracy (Shadmehr, Smith, & Krakauer, 2010; Ullsperger, Danielmeier, & Jocham, 2014). However, when external feedback is unavailable, individuals must rely on internal representations and neural mechanisms to regulate their actions. Without that feedback, even the most straightforward tasks, like cooking eggs, can quickly turn into a disaster, leaving us with a burnt mess.

In the lab, the role of feedback on force control has focused on vision and is commonly investigated using measures of isometric force control, for example, in experiments where participants are asked to maintain constant grip force at a target level (Poon, Chin-Cottongim, Coombes, Corcos, & Vaillancourt, 2012; Vaillancourt & Russell, 2002; Abolins & Latash, 2022; Baweja, Kennedy, Vu, Vaillancourt, & Christou, 2010; Menghi, Coricelli & Hickey, 2025). When visual performance feedback is provided, participants can adapt their grip force to better approximate the target. However, in the absence of feedback, the internal representation of the target force can degrade over time due to noise (Manohar et al., 2015; Slifkin & Newell, 1999), limited memory capacity (Poon et al., 2012; Vaillancourt & Russell, 2002), and the absence of error signals to guide correction (Abolins & Latash, 2022; Baweja et al., 2010). As a result, force output tends to drift, reflecting diminishing ability to accurately monitor and sustain the intended motor output based solely on internal cues.

The feedback loop between sensory input, performance monitoring, and motor adjustment is believed to rely on distinct and temporally coordinated neural processes. Sensory and perceptual systems support a dynamic update of internal models of action through interactions between somatosensory and motor cortices (Monaco, Fattori, Galletti, & Goodale, 2006; Noble, Eng, & Boyd, 2013; Sartin, Ranzini, Scarpazza, & Monaco, 2023). This ongoing exchange forms a closed-loop control system, allowing the brain to compare expected and actual outcomes in real time (Li, Sarma, Sejnowski, & Doyle, 2023). The dynamics of this loop appear to be reflected in oscillatory brain activity, particularly in theta (4–8 Hz) and beta (12–30 Hz) frequency bands, which are thought to orchestrate the continuous evaluation and adjustment of motor output. Brain activity in these frequency ranges has been closely associated with the coordination of performance monitoring (Kilavik, Zaepffel, Brovelli, MacKay, & Riehle, 2013), motor inhibition (Hannah & Aron, 2021; Logan & Cowan, 1984; Wessel & Anderson, 2023), and adaptive responses to errors (Little, Bonaiuto, Barnes, & Bestmann, 2019).

Theta oscillations (∼6Hz), which originate from the medial frontal cortex and supplementary motor areas, play a particularly crucial role in conflict detection and performance monitoring (Cavanagh & Frank, 2014; Cohen, 2014; Nachev, Kennard, & Husain, 2008). Theta is involved in both the detection of errors and in the orchestration of performance adjustments, creating a cyclical temporal reference that guides information processing and the optimization of goal-directed behavior. When an error is detected, beta oscillations, which appear to maintain the status quo and support ongoing motor actions (Baker, 2007; Engel & Fries, 2010), are reduced, allowing for the initiation of corrective actions and recalibration of performance (Chakarov et al., 2009; Duque, Greenhouse, Labruna, & Ivry, 2017; Huster, Enriquez-Geppert, Lavallee, Falkenstein, & Herrmann, 2013; Neuper & Pfurtscheller, 2001). When error correction is contingent on feedback, attentional alpha oscillations play an additional role, support the reallocation of attentional resources and facilitate the recalibration of motor actions in response to errors (Carp & Compton, 2009; Klimesch, 2012; Savoie, Thénault, Whittingstall, & Bernier, 2018; Wilken et al., 2023).

While much is known about neural responses to discrete errors, prior studies have predominantly used paradigms in which monitoring occurs only briefly or where corrections are triggered by isolated events (Holroyd & Coles, 2002; Pezzetta, Nicolardi, Tidoni, & Aglioti, 2018; Yeung, Botvinick, & Cohen, 2004). These one-shot corrections offer only a partial view of the brain’s capacity for dynamic error regulation. Real-world motor behavior often involves ongoing fluctuations in performance, where the magnitude of error can change moment to moment. In these situations, it is not a single, stable error that must be corrected, but rather a dynamically evolving discrepancy that requires continuous monitoring and adaptation. While many studies have linked error processing to well-established neural systems, the mechanisms that support ongoing adjustment during sustained performance are less clearly understood. Dynamic correction may involve processes such as sustained monitoring, integration of sensory feedback, and continuous performance recalibration, potentially engaging overlapping, but not identical, neural dynamics (Cohen, 2016; Wilken et al., 2023).

In order to explore this possibility, experimental designs must more closely approximate real-life sensorimotor demands and occur over real-life temporal intervals. With this in mind, the current study employs an isometric force control paradigm in combination with electroencephalography (EEG) to investigate the neural mechanisms supporting online motor correction. Participants were instructed to maintain a constant grip force at a target level under conditions where visual feedback about their performance was either continuously available or briefly available at the beginning of the trial (Early Feedback). This design allowed us to track how fluctuations in behavioral output correspond to dynamic changes in neural activity, offering a window into the closed-loop system of motor control. The early feedback condition was included as a control condition to compare how the system initially fails to recalibrate once that feedback is removed, providing insight into error recalibration versus performance decay.

By examining the temporal relationship between force deviations and EEG signals, we aimed to characterize the cortical processes that support rapid error detection and correction in real time. Our results show that when visual feedback was available, participants’ force output exhibited rhythmic fluctuations at ∼6 Hz, consistent with continuous online correction. These behavioral oscillations were accompanied by a dynamic pattern of EEG activity in the theta, beta bands and alpha bands, which tracked exertion and performance and predicted adjustments in motor output. In contrast, when feedback was absent, performance decayed linearly over time, and neural responses shifted, reflecting reduced engagement of real-time correction mechanisms.

## 2 Methods

### 2.1 Participants

Twenty-three participants (13 females, 10 males; mean age 24.2 years; range 20-28 years) gave informed consent before completing the experiment. The participants were all right-handed and naive to the purpose of the experiment. Two female participants were excluded from the analysis of the experiment because they commonly failed to respond, particularly in experimental conditions where feedback was not provided, resulting in a force error that was more than 3 standard deviations from the group mean. The experiment took approximately 2 hours to complete and all participants received a 20 euro reimbursement. All gave informed written consent and the study procedure was approved by the local institutional review board of the University of Trento.

### 2.2 Apparatus and Stimuli

Participants sat at approximately 60 cm from a computer monitor (VIEWPixx/EEG 22”; 1920x1080; 100 Hz) in a dimly illuminated room with their right hand lying over the table grasping a hand dynamometer. The dynamometer (HD-BTA Vernier) was used to record power grip force effort in Newtons (N) with an accuracy of `±`0.6 N. This dynamometer is a strain-gauge-based isometric force sensor which amplifies force and converts it into a voltage signal. The voltage signal was transferred to an Arduino Uno through Vernier interface shield hardware and subsequently to an acquisition computer. The force signal was sampled at 100 Hz. During the experiments, signals from this sensor were sent to MATLAB (The Mathworks Inc.) for visual real-time feedback of participant’s effort exertion. Feedback was updated at a frequency rate of 100 Hz.

Presentation of visual stimuli and acquisition of behavioral data was accomplished using PsychToolBox (Brainard, 1997) and custom MATLAB scripts. Before beginning the experiment, participants were requested to exert the most force they could on the dynamometer 3 times, each time for 3 s., with 10 s. of rest between each instance. The maximal voluntary contraction (MVC) was computed as the average of the highest peaks achieved in each of these trials. The trial sequence is illustrated in Fig. 1A. Each experimental trial began with a cue indicating the feedback condition, and then a target force appeared, which was randomly selected from 2 possibilities and calculated as a percentage of MVC (40% and 55%). Participants attempted to match this target force level with the hand dynamometer using a whole-hand power grip. For half of the trials, participants were presented with online visual feedback, for the other half they were presented with visual feedback for 1.5 seconds only then they had to rely on somatosensory inputs only.

**Figure 1:**
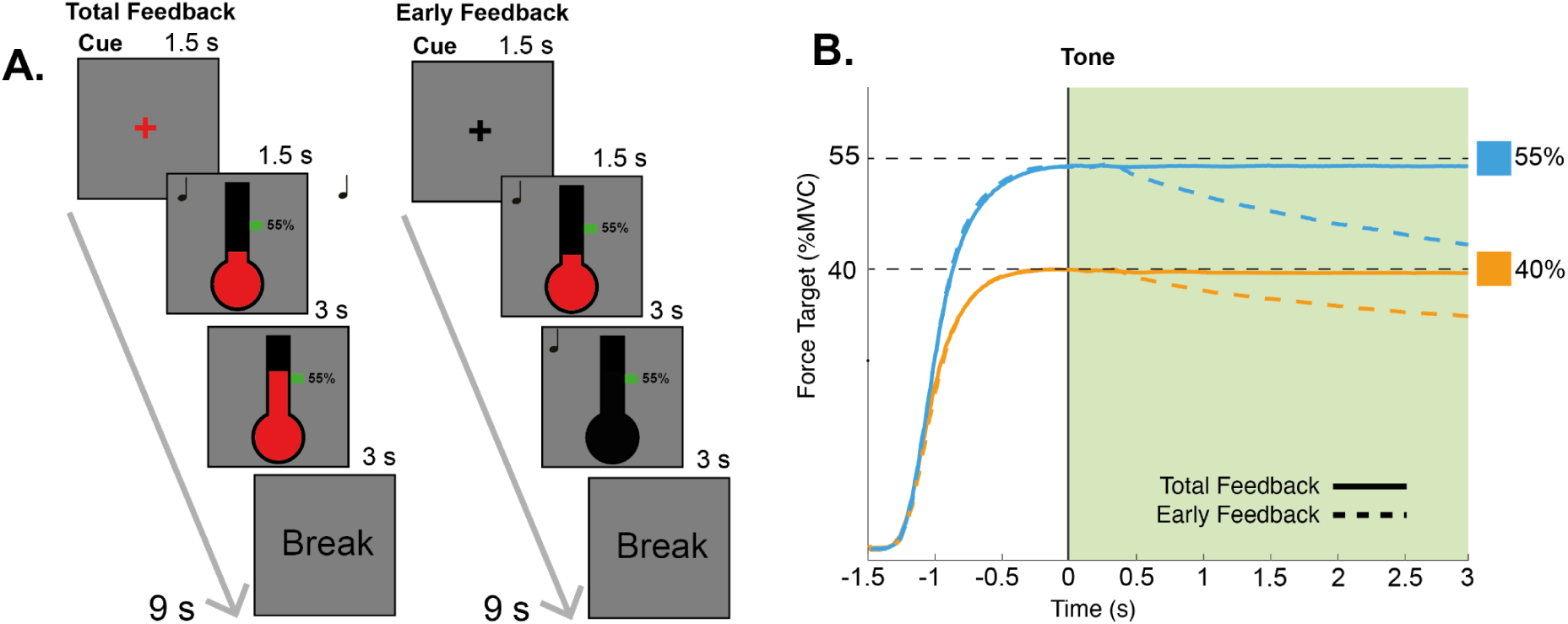
Panel A - Task schematics. Each trial began with the presentation of a fixation cross (1.5 s), which signaled the upcoming condition through its color (red or blue). An auditory cue then marked the start of the trial, triggering the appearance of the feedback display. If present, feedback was shown as a red fluid inside a stylized thermometer. The task lasted 4.5 seconds, consisting of a 1.5-second force estimation phase and a 3-second maintenance phase, each indicated by an auditory cue. This was followed by a 3-second relaxation period, making each trial 9 seconds in total. Participants received feedback throughout both force estimation and maintenance phases in the Total Feedback condition, while in the Early Feedback condition, feedback was provided only during the force estimation phase. At the end of each block, a message prompted participants to relax. **Panel B - Force estimation and maintenance.** This panel shows the average performance across participants in the total feedback and early feedback conditions and in the two different force targets.

When visual feedback was present, it took the form of a stylized black thermometer that was displayed at the center of an otherwise uniform dark grey background. The thermometer became increasingly red as force was exerted on the dynamometer. A green square on the thermometer indicated the target force output. When visual feedback was absent, the thermometer stayed on screen but the red ‘fluid’ was not presented. Task performance lasted 4.5 s. and began with an auditory tone indicating the beginning of a 1.5 s. force estimation period, during which participants were to adjust the force to the target value. A tone subsequently indicated the beginning of a 3 s. maintenance period and a final tone indicated the end of the trial. The experiment was composed of 15 practice trials followed by 160 experimental trials divided into 8 blocks, with breaks between blocks.

### 2.3 EEG Preprocessing

EEG was recorded at 1 kHz from 64 Ag/AgCl electrodes mounted in an elastic cap (10/20 montage) using the BrainAmp-DC system and Brain Vision Recorder software (Brain Products). Additional electrodes were placed at the left and right mastoids and 1 cm lateral to the outer canthus of the left eye. All electrodes were referenced during the recording to the right mastoid. Electrode impedances were kept below 10kΩ. Preprocessing was carried out using Fieldtrip Toolbox for MATLAB (Oostenveld, Fries, Maris, & Schoffelen, 2011). Continuous data were re-referenced to the common average and highpass filtered at 0.1 Hz with a zero-phase sixth-order Butterworth filter. The data were epoched from 2.5 seconds before the beginning of the maintenance period (See Fig.1 panel B) to 3.5 seconds following it. We visually inspected these epochs to remove trials containing muscle activity and electrical artifacts and to identify bad channels, which were interpolated to the weighted average of the neighboring electrodes. Fast Independent Component Analysis (fastICA) (Hyvärinen, 1999) was then performed on the epoched data. ICA components representing eye blinks, eye movements and sustained high-frequency noise were visually identified and removed from the data. No trials were discarded during this procedure. EEG epochs were low-pass filtered with a cut-off of 100Hz and notch filtered at 50Hz. Finally, we visually re-inspected the epochs to ensure no residual artifacts. Rejected trials were excluded from all further analyses.

### 2.4 Dynamometer spectral analysis

To evaluate the presence of periodic phenomena during the force maintenance phase, we applied a series of preprocessing steps of the signal over time recorded from the dynamometer. We first selected the 3 seconds corresponding to the maintenance phase. For each participant and each trial, the signal was first normalized in percentage of the MVC, detrended (first order) and further z-scored along the time dimension. In order to separate fractal and periodic components from the signal in the frequency domain, we adopted the IRASA algorithm (Wen & Liu, 2016) as implemented in fieldtrip. The periodic component was obtained by dividing the spectra by the fractal component, yielding, for each participant, one spectral profile of the oscillatory component contained in the dynamometer signal in each combination of force (40% vs 55%) and feedback (present vs absent).

We wanted to further evaluate the presence of oscillatory phenomena arising specifically as a function of feedback. For this reason, in each frequency bin from 2 to 20 Hz we computed a Linear Mixed Effect Model (LMEM, implemented in Python <u>statsmodels</u> 0.14.4) with force and feedback as main effects and subjects as a grouping variable to account for repeated measures:

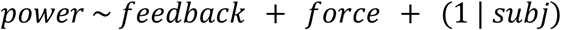

The density of frequency bins and the extension of the window of interest in the frequency domain implied a high number of LMEM models tested, which made it necessary to account for multiple comparisons. This was achieved by applying a cluster permutation approach (Maris & Oostenveld, 2007), where we first defined clusters by selecting the contiguous, absolute z values exceeding the critical threshold of the 95th percentile of a normal distribution. All the contiguous values in each cluster were then summed together. Then, for 1500 iterations, the experimental labels were shuffled in each participant, creating in this way a set of surrogate datasets. In each of the shuffled datasets, we repeated the computation of the LMEM in each frequency bin, as well as the clustering, and in each permutation, we collected the cluster statistic with the highest absolute value into a distribution of cluster statistics under the null hypothesis. Finally, we compared the cluster statistics obtained by our tests on the empirical data to the distribution of cluster statistics under the null hypothesis, and we rejected the null hypothesis if a cluster statistic was below the 2.5th or above the 97.5th percentiles of the permutations.

### 2.5 EEG-Dynamometer analysis

To identify alignment between behavioral and EEG datasets, we downsampled the EEG data to 100 Hz to match the sample rate of the behavioral data, and focused on results observed in both signals during the maintenance period. Statistical analysis relied on a cluster-based permutation approach. Statistical significance was assessed non-parametrically at the group level using a cluster-based permutation approach with a cluster-forming threshold of p < 0.05 (two-tailed), and a corrected significance level of p < 0.05 (two-tailed) (Maris & Oostenveld, 2007). Condition labels were randomly permuted 1000 times.

## 3 Results

Participants performed a force production task using a hand dynamometer, where in each experimental trial they were required to match one of two possible force targets under different feedback conditions (see Fig. 1). In the Total Feedback condition, participants were visually provided with online performance feedback throughout each 5 s trial. In the Early Feedback condition, feedback was presented for only the first 1.5 seconds. As illustrated in Figure 1B, participants were able to accurately maintain force during the total feedback condition, but force production decayed when only early feedback was provided.

To investigate the temporal dynamics of continuous, visually-guided error correction, we conducted two streams of data analysis. In the analysis of behavior, we compared participant performance between the total feedback and early feedback conditions by applying spectral analysis. This allowed us to identify potential oscillatory patterns in force generation during action maintenance. In a subsequent analytic stream, we used cross-correlation to investigate the relationship between EEG (in both the time and frequency domains) and two behavioral measures: force production and the degree of error, defined as deviation from the target force.

### 3.1 Behavioral Results ***-*** Spectral Analysis

We began by exploring the oscillatory structure of behaviour in our force target conditions (40% or 55% of Maximum Voluntary Contraction; MVC). Results showed that force generation had a ∼6 Hz oscillatory component that emerged only when performance feedback was available (See Fig.2A). In the raw data, this ∼6 Hz effect is couched in a strong 1/f structure, with power decreasing as a function of frequency. To isolate our effect from this overall pattern, we removed the aperiodic component of this signal (Fig. 2B; see *Dynamometer spectral analysis* in *Methods*).

**Figure 2:**
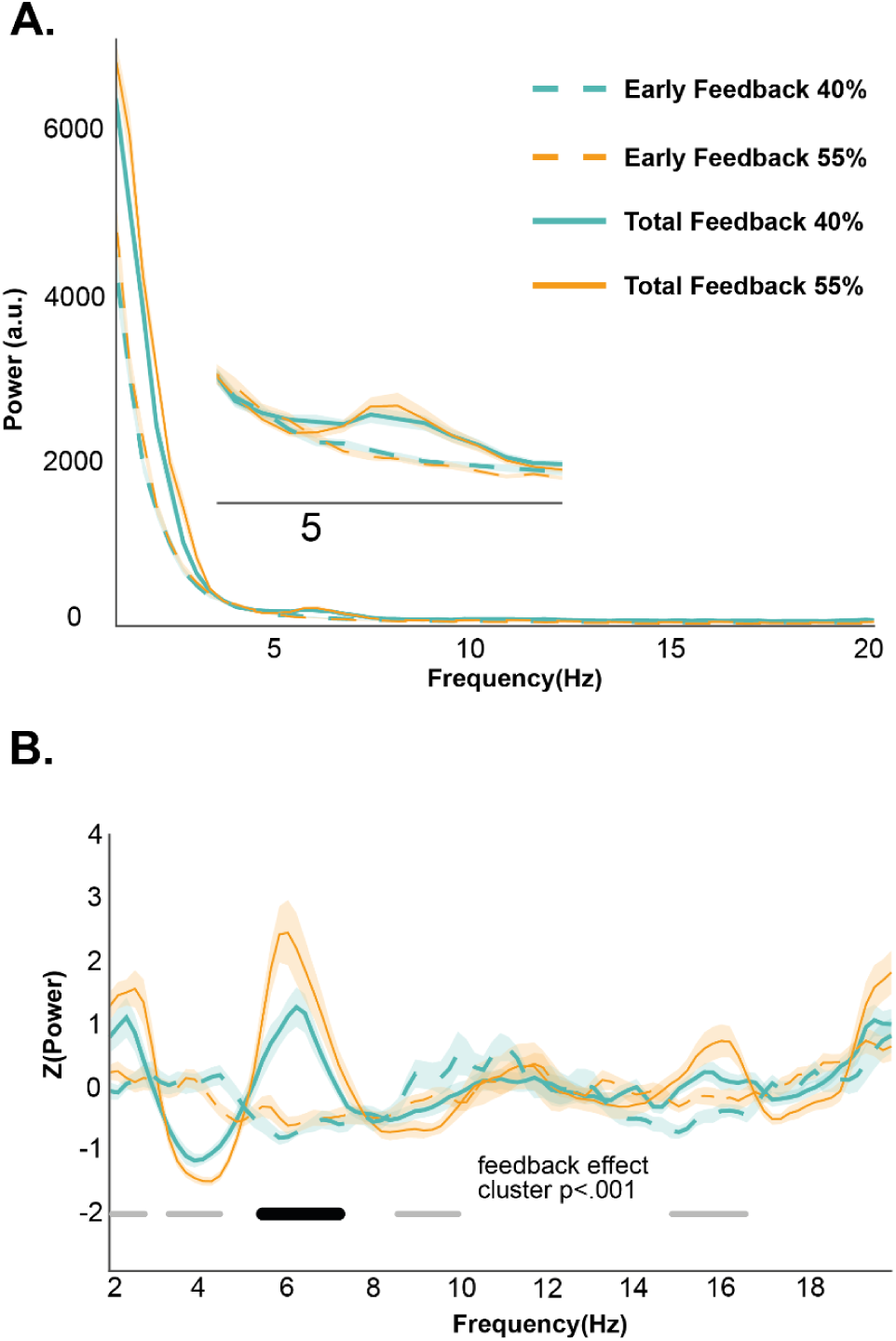
Spectral and linear analyses of force output across feedback conditions. **(A)** Power spectral density of the force signal across the different feedback and target conditions. Feedback condition shows a 6Hz rhythmic fluctuation during the exertion. This peak is absent in the no-feedback condition. This is more evident in the sub-panel, which reflects a magnification of the 4 - 8 Hz portion of the larger panel. **(B)** Spectral density after removing the aperiodic 1/f component from the signal.

The effect emerges in this cleaned plot between 5.27 Hz and 7.42 Hz, with congruent peak frequencies in the two force conditions (40%=6.25 Hz; 55%=6.05 Hz). To statistically test this effect, we used linear mixed modelling with feedback presence as a predictor of dynamometer’s power in each frequency bin. We focused our analyses over the range between 2 and 20 Hz to evade both slow trends that could not clearly be associated with oscillatory patterns in the dynamometer signal, as well as faster oscillations that were not the target of our hypotheses. Results were cluster-corrected for multiple comparisons (see *Methods*). This analysis confirmed the presence of a significant effect of feedback over the interval between 5.27 Hz and 7.42 Hz (cluster p <.001).

Our approach also revealed significant effects of feedback in the delta (3.32-4.88 Hz) and in the mu ranges (8.59-9.96 Hz), as well as other short-duration effects at low delta range (1.95-2.73 Hz) and low beta range (17.18-18.16 Hz). These effects tended to emerge in periods of low frequency power in both feedback conditions, or in intervals where the separation of periodic and aperiodic effects is problematic, making interpretation difficult. With this in mind, we have focused analysis on effects in the theta range, where strong peaks in oscillatory power emerge.

### 3.2 EEG Results

To explore the relationship between behavioral performance and EEG activity during the maintenance period, we approached our results in three separate ways. First, we examined the cross-correlation between overall behavioral performance and the broadband EEG signal, allowing us to identify general patterns of neural activity linked to task performance. Next, we focused on the relationship between behavioral error, defined as the absolute distance from the target, and EEG. This included both broadband EEG and power within the theta (4-8Hz), alpha (8-12Hz), and beta (12-30Hz) frequency bands. Finally, we calculated the cross-correlation between the behavioral oscillatory pattern and power in the theta, alpha and beta frequency bands described above.

#### 2.2.1 Cross-correlation between performance and broadband EEG

For each trial, we computed the cross-correlation between behavioral exertion and EEG. The two signals were shifted relative to each other in time units of 0.01 seconds, to a maximum lag of 1 s, with correlation computed at each step. In the total feedback condition, we found that the behavioral data were positively correlated with the EEG with significant clusters at positive time lags (-20 to 350 ms and 300 to 600 ms). This indicates a relationship in which dynamometer activity predicted the magnitude of the subsequent EEG signal. These effects emerge in EEG recorded from central electrodes, consistent with generators in the somatosensory cortex (see Fig. 3 Panel A). In the early feedback condition, we observed a prolonged negative correlation between the EEG signals and force exertion, spanning from positive to negative time lags across the entire scalp (see Fig. 3 Panel B). This appears to reflect the greater decay in behavioural performance in this condition. Without continuous feedback, participants likely relied more on internal representations of the target force, which decayed over time. The widespread nature of the correlation suggests that, in the absence of external feedback, neural activity tracks a general decline in performance rather than discrete corrective adjustments. Additionally, we identified a positive correlation at positive time lags, specifically from *250 ms* to *1 s*, localized in the posterior region of the scalp. We interpret this as reflecting an evaluation of force exertion in the absence of online feedback, potentially supporting recalibration based on internal sensory predictions. No differences were found between the two force targets in the total feedback or the early feedback conditions.

**Figure 3:**
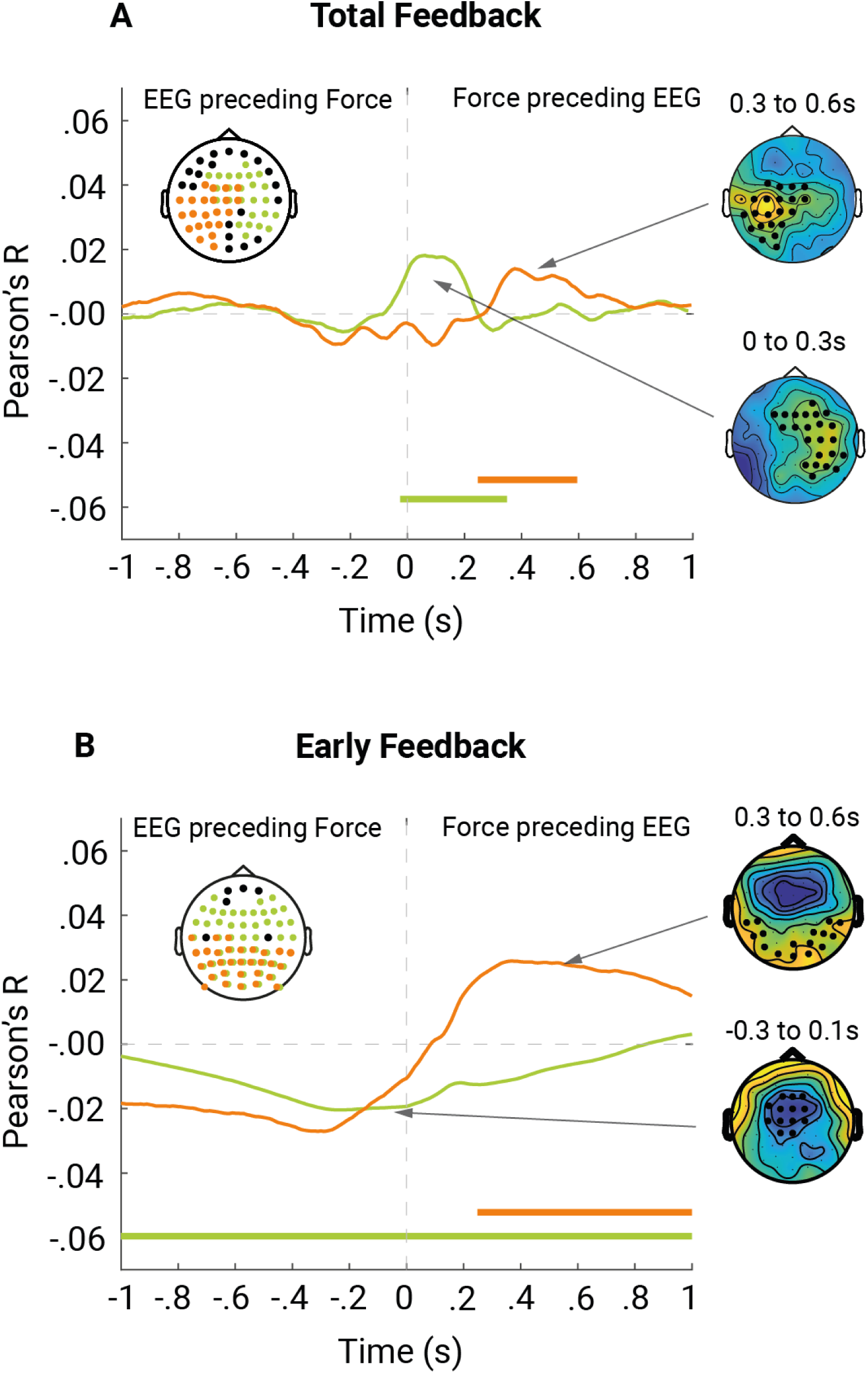
Cross-correlation between behavioral performance and broadband EEG signal in the Total Feedback **(A)** and Early Feedback **(B)** conditions. Positive and negative lags indicate whether EEG activity precedes or follows behavioral performance. Significant clusters are highlighted in green and orange, with the corresponding time intervals marked with the same color code at the bottom. The white topoplot identifies the sensors contributing to these significant clusters, while additional topographical plots illustrate the spatial distribution of EEG activity at key peaks within each condition and the relative significant channels highlighted.

#### 2.2.2 Cross-correlation between behavioral error and broadband EEG

When examining the relationship between EEG broadband signals and absolute error from the target, we restricted our analysis to the total feedback condition only, as performance in the early feedback condition consistently decayed over time, making error from the target less meaningful. For each trial, we computed the cross-correlation between behavioral error and EEG in 0.1 s steps and a maximum lag of 1 second. Two clusters emerged with early positive lags (Figure 4A), reflecting a situation where dynamometer activity predicted the amplitude of the subsequent EEG signal. In the earlier cluster, we observed a positive correlation, while the later cluster was defined by negative correlation, such that a higher absolute distance from the target (ie. error) was associated with reduced EEG amplitude. The negative cluster observed at -0.50 ms to 940 ms spanned from early posterior to later central electrodes. The topography of this effect is consistent with generators in medial prefrontal and anterior cingulate cortex, brain areas known to respond to error commission. The positive cluster at time lag *-140 ms* to *410 ms* appears to reflect a generator in the frontal cortex. This may indicate anticipatory processes or preparatory adjustments preceding changes in behavioral output, potentially reflecting proactive monitoring and control mechanisms.

**Figure 4:**
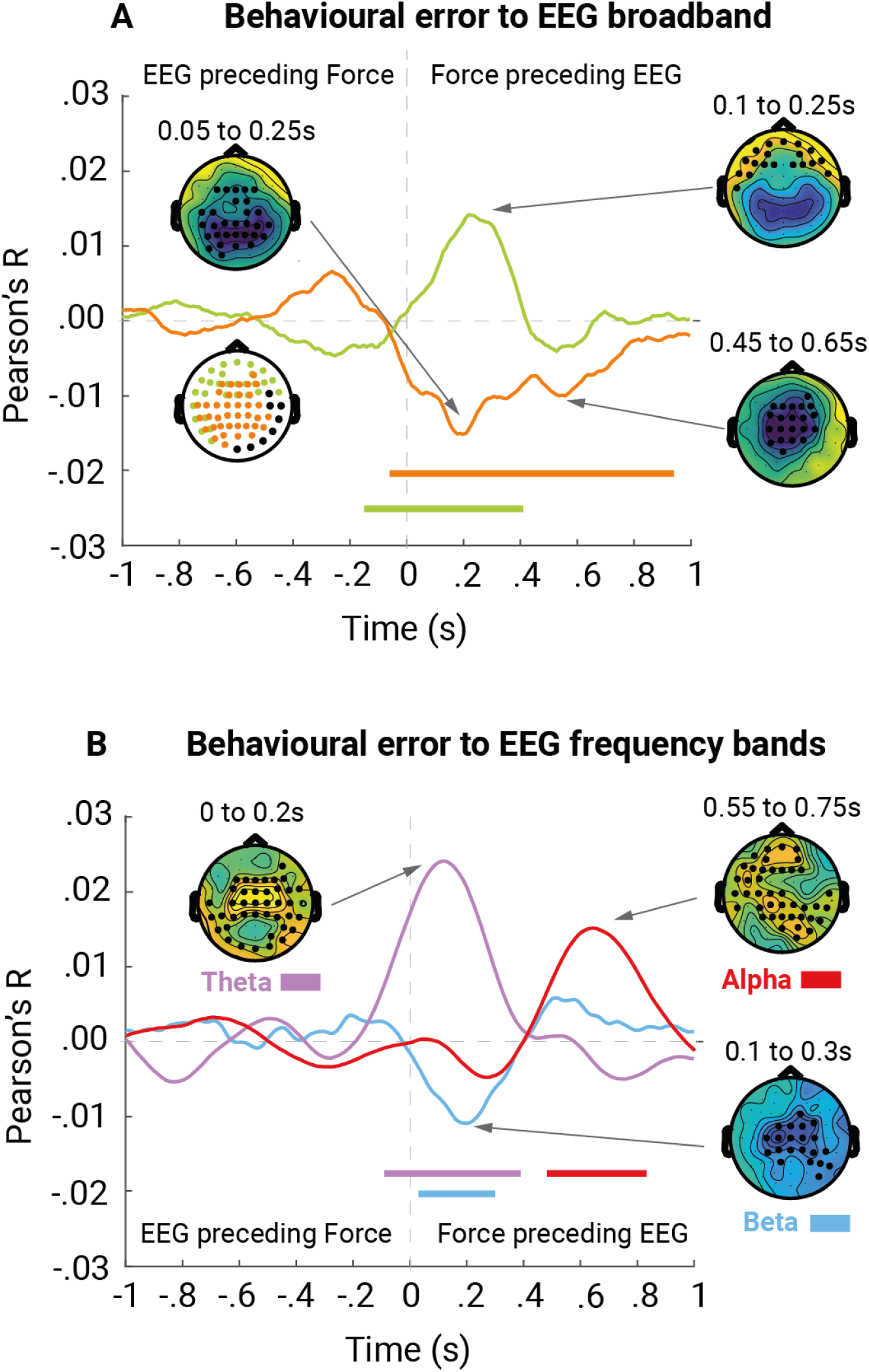
Cross-correlation between behavioral error from the target and broadband (A) or frequency band-specific (B) EEG signal in the Total Feedback condition. Positive and negative lags indicate whether EEG activity precedes or follows behavioral performance. Significant clusters are highlighted in different colors, with the corresponding time intervals marked with the same color code on top. The white topoplot identifies the sensors contributing to these significant clusters, while additional topographical plots illustrate the spatial distribution of EEG activity at key peaks within each condition, with significant channels highlighted.

#### 2.2.3 Cross-correlation between behavioral error and EEG power in theta, alpha and beta bands

We computed the cross-correlation between absolute behavioral error and EEG power in theta, alpha and beta frequency bands in the total feedback condition, allowing for a maximum lag of 1 second. EEG power was estimated using the Hilbert transform to obtain the instantaneous power in the theta (4–8 Hz), alpha (8–12 Hz), and beta (13–30 Hz) frequency bands. Our analysis revealed a significant positive cluster in each frequency band (with three independent analyses), all occurring at positive lags (See Figure 4 Panel B). This indicates that fluctuations in absolute behavioral error preceded corresponding changes in EEG power. The first significant cluster appeared in the theta band, suggesting an early neural response to behavioral error. Shortly thereafter, a significant cluster emerged in the beta band, overlapping temporally with theta but peaking slightly later. Finally, a significant alpha cluster was observed, emerging as the theta and beta effects began to decay. This temporal progression suggests a sequential recruitment of oscillatory activity, potentially reflecting distinct stages of error processing, motor correction, and feedback monitoring.

#### 2.2.4 Cross-correlation between behavioral oscillatory power and EEG power in theta, alpha and beta bands

Finally, we examined the cross-correlation between the power of behavioral oscillations in the total feedback condition and EEG power, allowing for a maximum lag of 1 second. Behavioral oscillations were identified in the 5.5–7.2 Hz range, according to behavioral results (see section 4.1), and their instantaneous power was again extracted using the Hilbert transform. Similarly, EEG power was identified in the theta (4–8 Hz), alpha (8–12 Hz), and beta (12–30 Hz) frequency bands. Our analysis revealed a single significant positive cluster in the theta frequency band, occurring at a positive time lag (See Figure 5). This indicates that fluctuations in the power of ∼6 Hz oscillations in force generation precede corresponding changes in theta power in the EEG. No significant clusters were observed in the alpha or beta bands. This result suggests a role of theta oscillations in processing force-related feedback.

**Figure 5:**
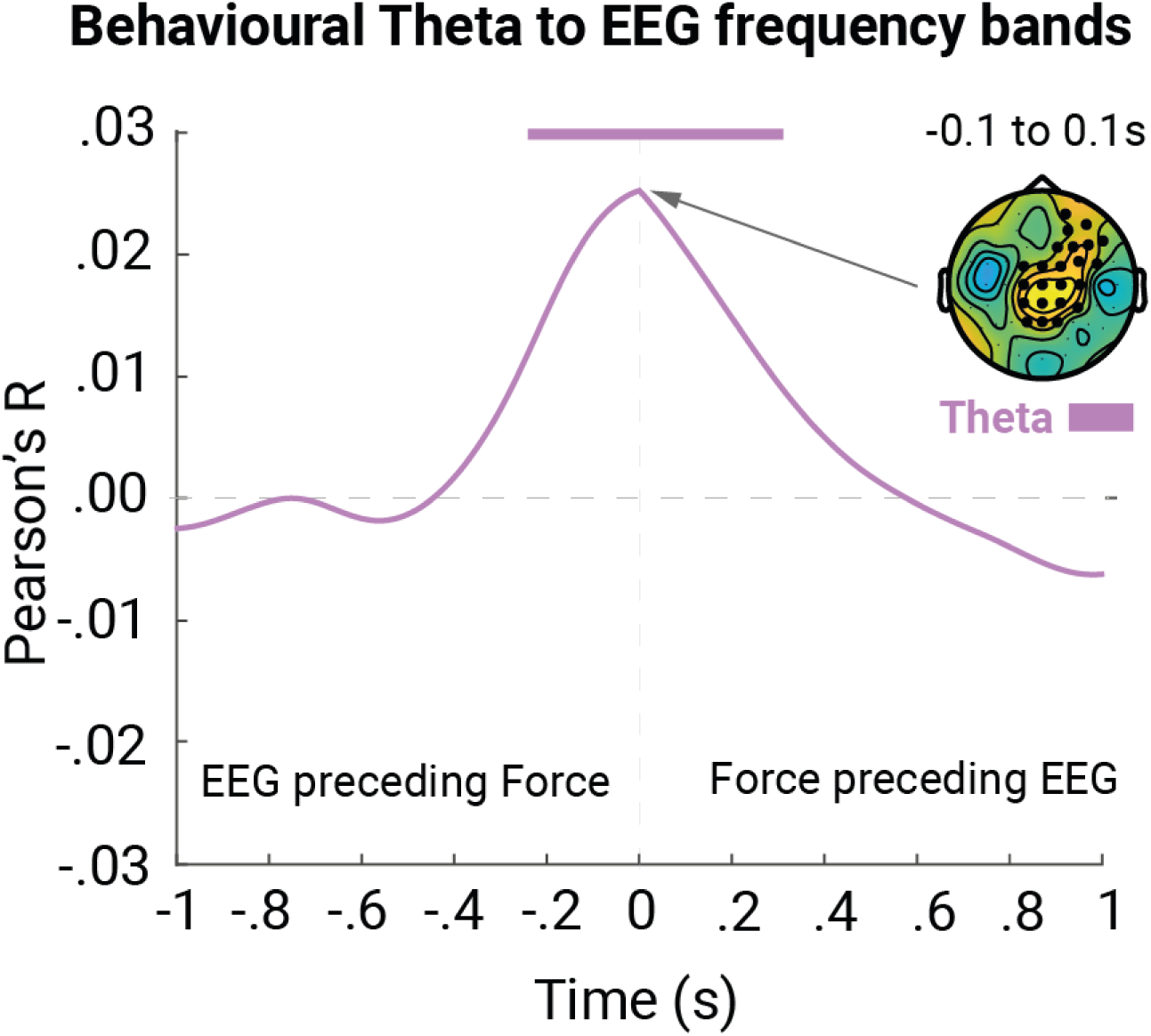
Cross-correlation between behavioral and EEG power in the theta band in the Total Feedback condition. Positive and negative lags indicate whether EEG activity precedes or follows behavioral performance. The time course of the significant cluster is presented, with the corresponding time intervals marked with the same color code on top. The topographical plots illustrates the spatial distribution of EEG activity at the key peak, significant channels are highlighted.

## 4 Discussion

We investigated the temporal dynamics of visually guided error correction during a task that required participants to continuously adjust hand force to match a target. Behaviorally, we found that continuous visual feedback facilitated error correction, helping participants maintain their force output close to the target. In the presence of feedback, performance exhibited a peak of performance fluctuation centered at approximately 6 Hz, suggesting that correction was largely implemented at this rate. The emergence of this specific frequency band is intriguing, as oscillations in this theta range have been observed across a variety of everyday behaviors requiring ongoing monitoring and fine adjustments, such as speech production, planning, typing and visuo–motor coordination (Kaplan et al., 2020; Ten Oever, Titone, Te Rietmolen, & Martin, 2024; Vallbo & Wessberg, 1993; Yamaguchi, Crump, & Logan, 2013). This has led to the proposal that theta oscillations may serve as an internal metronome, facilitating the continuous evaluation and adjustment of motor output in response to performance errors (Cohen, 2014). Theta oscillations in behaviour emerged in our results only in the total feedback condition. In the early feedback condition, where visual feedback was discontinued early in the trial, force output instead gradually decayed without evidence of correction (See also Abolins & Latash, 2022; Vaillancourt, Thulborn, & Corcos, 2003; Limonta, Rampichini, Cè, & Esposito, 2015).

We examined the relationship between ongoing behaviour and ongoing brain activity using a number of techniques. We began by examining the cross-correlation between force exertion and EEG amplitude as observed in the total feedback condition, finding a positive relationship in EEG electrodes located over somatosensory cortex, suggesting a strong sensorimotor coupling during force adjustments. In contrast, in the early feedback condition, exertion was associated with activity in central and occipital channels. This pattern may reflect the gradual decline in performance over time: without ongoing visual feedback, neural activity appears to track the decay of force rather than actively signaling or correcting for errors

Examining the correlation between broadband EEG activity and absolute error from the target, we found that larger errors were positively correlated with activity in frontal channels and negatively correlated with activity in central channels. The positive correlation in frontal regions may reflect increased engagement in performance monitoring, consistent with the role of the medial frontal cortex in detecting deviations from expected outcomes (Badre & Wagner, 2005; Luu, Flaisch, & Tucker, 2000; Ridderinkhof, Ullsperger, Crone, & Nieuwenhuis, 2004). The negative correlation in central regions could be linked to error-related negativity, a well-documented neural response to performance errors (Trujillo & Allen, 2007; Wessel, 2012).

Next, we examined the cross-correlation between absolute error and EEG instantaneous power in the theta (4-8 Hz), alpha (8-12 Hz), and beta (12-30 Hz) frequency bands. Behavioral error was initially positively correlated with occipito-central theta, followed shortly by a negative correlation with central beta, and finally, a positive correlation with fronto-central alpha. We interpret the early theta-band correlation as reflecting activity in medial frontal regions responsible for cognitive control and error processing (Cavanagh & Frank, 2014; Cohen, 2014; Nachev et al., 2008). The subsequent beta suppression is likely to reflect the initiation of motor responses in order to correct behaviour, and specifically to discontinue ongoing ‘status quo’ performance (Engel & Fries, 2010). This interpretation is consistent with a recent study demonstrating a decrease in beta power during muscle contraction during reaching (Matta, Baurès, Duclay, & Alamia, 2025). Finally, subsequent frontal and occipital alpha modulation is consistent with the idea that participants adapt attentional control for visuomotor processing to monitor their performance adjustment (Rilk, Soekadar, Sauseng, & Plewnia, 2011). This is in line with existing results suggesting connectivity in alpha and beta frequency bands underlies this kind of monitoring in visuomotor tasks (Hipp, Engel, & Siegel, 2011; Palva & Palva, 2007; Zhang, Chavarriaga, & del R Millán, 2015). This temporal sequence suggests an error-processing loop, where errors first engage cognitive control (theta), followed by correction of motor activity (beta), and finally, a recalibration of attentional and motor networks (alpha) to optimize subsequent performance.

Finally, we cross-correlated the instantaneous power of the behavioral exertion signal in the significant frequency band with the instantaneous power of EEG activity in the theta, alpha, and beta frequency bands. We found a significant correlation with central theta, coherent with a role for theta oscillations in sensorimotor coordination (Cohen, 2016). This finding indicates that fluctuations in exertion are dynamically linked to neural processes involved in continuous feedback processing and error correction mechanisms.

In summary, our findings reveal the neural dynamics behind visually guided force control and error correction. Feedback processing involved a dynamic interplay of oscillatory activity, with theta, gamma and alpha bands reflecting different stages of performance monitoring and motor adaptation. Together, these results provide insight into the neural dynamics of motor control and suggest that theta oscillations may serve as a bridge between sensory feedback and motor execution, facilitating adaptive behavior during completion of ongoing, continuous tasks.

## Acknowledgments

This project was supported by the Autonomous Province of Trento, Italy (call ’Grandi Progetti 2012,’ project ’Characterizing and improving brain mechanisms of attention – ATTEND’). C.H. is further supported by the European Research Council under the European Union Horizon 2020 Research and Innovation Program (804360).

